# Crossmodal visual predictions elicit spatially specific early visual cortex activity but later than real visual stimuli

**DOI:** 10.1101/2022.12.14.520404

**Authors:** Liesa Stange, José P. Ossandón, Brigitte Röder

## Abstract

Previous studies have indicated that crossmodal visual predictions are instrumental in controlling early visual cortex activity. The exact time course and spatial precision of such crossmodal top-down influences on visual cortex have been unknown. In the present study, participants were exposed to audio-visual combinations comprising one of two sounds and a Gabor patch either in the top left or in the bottom right visual field. Event related potentials (ERP) were recorded to these frequent crossmodal combinations (Standards) as well as to trials in which the visual stimulus was omitted (Omissions) or the visual and auditory stimuli were recombined (Deviants). Standards and Deviants elicited an ERP between 50 and 100 ms of opposite polarity known as a C1 effect commonly associated with retinotopic processing in early visual cortex. In contrast, a C1 effect was not observed in Omission trials. Spatially specific Omission and Mismatch effects (Deviants minus Standards) started only later with a latency of 230 ms and 170 ms, respectively.

These results suggest that crossmodal visual predictions control visual cortex activity in a spatially specific manner. However, visual predictions do not elicit neural responses that mimic stimulus-driven activity but rather seem to affect early visual cortex via distinct neural mechanisms.

## Introduction

Expectations about forthcoming sensory input in a continuously changing environment are essential for efficient behavior: Sensory predictions have been reported to lower detection thresholds and increase processing speed for upcoming stimuli (de Lange, Rahnev, Donner, & Lau, 2013; Den Ouden, Friston, Daw, McIntosh, & Stephan, 2008; Puri & Wojciulik, 2008; Puri, Wojciulik, & Ranganath, 2009). In a multisensory world, sensory predictions are possible based on input of another sensory modality. For example, auditory stimuli preceding visual events are capable of changing the perception of the latter (Feng, Störmer, Martinez, McDonald, & Hillyard, 2014).

The predictive coding framework proposes that top-down predictions and bottom-up sensory input are compared at sensory processing stages; if predictions do not match the actual sensory input, this framework assumes that an error signal is generated which travels downstream to update representations about the world (Bastos et al., 2012; de Lange, Heilbron, & Kok, 2018). In line with these assumptions, both brain imaging (Alink, Schwiedrzik, Kohler, Singer, & Muckli, 2010; Den Ouden et al., 2008) and electrophysiological studies (Todorovic & Lange, 2012; Wacongne et al., 2011) have revealed enhanced neural responses to unexpected compared to expected sensory events, which were interpreted as error signals.

In order to isolate the neural signature of the predictive signal the response to expected but omitted stimuli has been investigated. For example, functional magnetic resonance imaging (fMRI) studies have confirmed that omitted visual stimuli modulate the primary visual cortex in a retinotopic specific manner (Kok, Failing, & Lange, 2014; Muckli et al., 2015). These results as well as analogous results in the auditory system (Todorovic, van Ede, Maris, & Lange, 2011) converge to the hypothesis that predictive signals are sent upstream and that they alter activity of early sensory cortex in a stimulus specific manner. However, yet the precise timing of how sensory predictions unfold their influence have been unknown because the time resolution of brain imaging techniques such as fMRI is insufficient to dissect visual cortex activity within the first 500 ms of the omitted stimulus. Thus, even if early visual cortex is well known to be subject of top-down control, e.g. based on crossmodal predictions, it is yet unclear when in time such a modulation emerges and when spatially selective error signals are generated. Some authors have suggested that predictions cause sensory cortex activity that mimics the sensory-driven activity. Thus, it is assumed that top-down and bottom-up generated neural activity are indistinguishable in early sensory cortex (Bendixen, Schröger, & Winkler, 2009). Testing this proposal requires an experimental protocol that allows tracking neural activity in early visual cortex with a millisecond resolution within the first few hundred milliseconds of stimulus processing.

The present event-related potential (ERP) study made use of a visual ERP named the C1 effect. The C1 comprises a polarity reversal of the ERPs between 50 and 100 ms for upper vs. lower visual field stimulation. This polarity reversal is considered to indicate activity of opposing banks of the calcarine sulcus and thus to reflect retinotopic processing in early, likely primary, visual cortex (Di Russo, Martínez, Sereno, Pitzalis, & Hillyard, 2001).

We associated the location of two circular gratings, one presented in the top left (V_top left_), and the other presented in the bottom right (V_bottom right_) quadrant of the visual field with two sounds differing in pitch. Consecutively, ERPs were recorded to these frequent learned associations (“Standards”), to rare trials in which the visual stimulus occurred at the unexpected location (“Deviants”) and rare trials in which the visual stimulus was not presented at all (“Omissions”).

We predicted to find a C1 effect for both Standards and Deviants. If crossmodal predictions mimic stimulus-driven activity in visual cortex, that is, neural activity, which is indistinguishable in time and space from stimulus-driven activity, we would expect to observe a typical C1 effect to Omissions in addition to often observed late omission effects (>200 ms). Moreover, we expected to find a typical visual mismatch effect to Deviant stimuli compared to Standard stimuli. If crossmodal predictions modulate but do not generate stimulus-driven activity in early visual cortex, we would predict a modulation of the Mismatch effect in the C1 and following time epochs by the visual prediction.

## Methods & Materials

### Subjects

Forty-five students of the University of Hamburg participated in the study. They all reported normal or corrected to normal vision, normal hearing and no history of psychiatric or neurological disorders. Data of two participants were excluded because of a too noisy electroencephalogram (EEG) (see below in section: EEG recording and preprocessing). Two other participants’ data sets were disregarded because data were missing for one condition, and data of one additional participant were removed due to extensive blinking. The remaining forty participants had an average age of 23.8 years (range: 18 to 45 years, SD = 5.8, 27 females, 5 left-handed). The local ethics board of the Faculty of Psychology and Movement Science of the University of Hamburg (Germany) had approved the study (No. 2018_190). All participants gave informed consent and received course credit as compensation.

### Stimuli and Apparatus

All stimuli were generated using the Psychophysics Toolbox for MATLAB (Brainard, 1997; Kleiner, Brainard, & Pelli, 2007). The visual stimuli were presented with a 23.5’’ Eizo FG2421 LCD monitor (Ishikawa, Japan) with a refresh-rate of 120 Hz. They consisted of full contrast circular grating patterns subtending an angle of 2.5°. The grating patterns were black and white horizontally oriented stripes with a spatial frequency of 2 cycles/°. In addition, some grating patterns were presented with vertically oriented stripes, which served either alone or in combination with an auditory stimulus as behavioral targets. The grating stimuli were presented one at a time for 500 ms. They were positioned in either the top left or bottom right visual quadrant at an eccentricity of 4°. The grating stimuli in the top quadrant were presented at an angle of 25° from the center (V_top left_), and the grating stimuli in the lower quadrant were displayed at an angle of 45° (V_bottom right_) (Figure 1A) to best target the opposing banks of the calcarine sulcus (Sourav, Bottari, Kekunnaya, & Röder, 2018). The auditory stimuli consisted of two, easily to differentiate sinusoidal tones (A1 and A2 at 1000 Hz and 400 Hz, respectively) with a duration of 1.25 s (including 83 ms linear rise and fall envelopes) presented at 70 dB(A) (as measured at the ear level of participants) from two speakers centrally positioned under the screen. In the auditory run, these tones were additionally presented with a 150 ms long gap inserted 750 ms post onset; these stimuli severed as behavioral targets (see below). During the whole experiment, participants were asked to indicate targets by pressing a Buddy Button (AbleNet, Inc., Minneapolis, United States of America) operated by their dominant hand.

**Figure 1.**
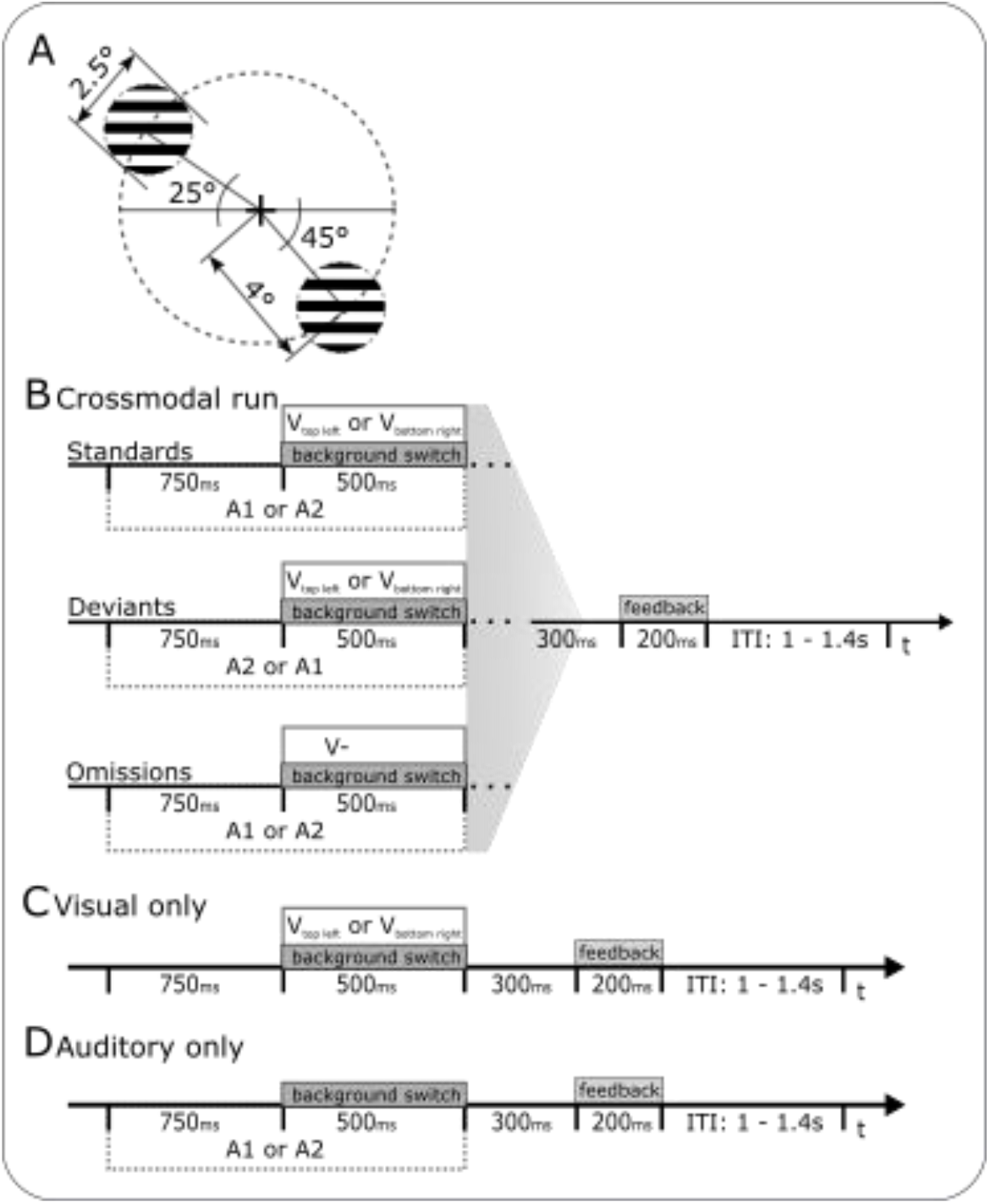
Visual stimulation and experimental protocol. (A) The visual stimuli comprised full contrast circular grating patterns, which were presented in either the top left or the bottom right quadrant, arranged such that they best hit opposite banks of the calcarine sulcus. (B) Trial structure of the crossmodal run separately illustrated for each of the three possible conditions from top to bottom: Standards, Deviants and Omissions. V_top left_= grating in the top left quadrant; V_bottom right_ = grating stimulus bottom right quadrant; A1=1000 Hz sound, A2= 400 Hz sound (assignment counterbalanced across participants); in each trial either V_top left_ or V_bottom right_ and either A1 or A2 were presented or no grating stimulus was presented at all (V-, Omissions). (C) Trial structure of the Visual only trials. (D) Trial structure of the Auditory only trials.

To ensure precise timing of the stimuli with respect to the event-trigger signal stored with the EEG file, a photodiode connected to a custom-made Arduino microcontroller (Banzi & Shiloh, 2014) was installed. A fixed delay of 23.74 ms (SD = 1.88) between the trigger event mark and the visual presentation was observed which was removed during the pre-processing of the EEG data.

### Design

#### Crossmodal run

The trial structure was the same for all conditions (Figure 1B): Trials started with the simultaneous presentation of a white central fixation cross (size 0.25°) on a black background and the onset of the auditory stimulus. The fixation cross remained visible until the end of a trial. After 750 ms the background was inverted from black to gray (called ‘background switch’ in the following) and simultaneously a grating stimulus was presented at one of the two possible locations. The grating and auditory stimulus were extinguished after 500 ms and the background changed back to black.

There were two frequent combinations of a sound and a grating location (termed “Standards”: A1V_top left_/A2V_bottom right_), which tone and grating location were paired was counterbalanced across subjects. Standards were presented in 70% of the trials. In 4% of the trials, the sounds and grating locations were recombined (called “Deviants”: A1V_bottom right_/A2V_top left_). In both conditions, gratings were presented with vertically oriented stripes. In 18% of the trials, the grating stimulus was omitted (called “Omissions” in the following: A1V-/A2V-) (see Table 1 for a summary of all conditions). Since a background switch occurred in Omission trials too, the time point of the omission was obvious to the subjects. In fact, the background switch was introduced to reduce temporal uncertainty in omission trials.

**Table 1.**
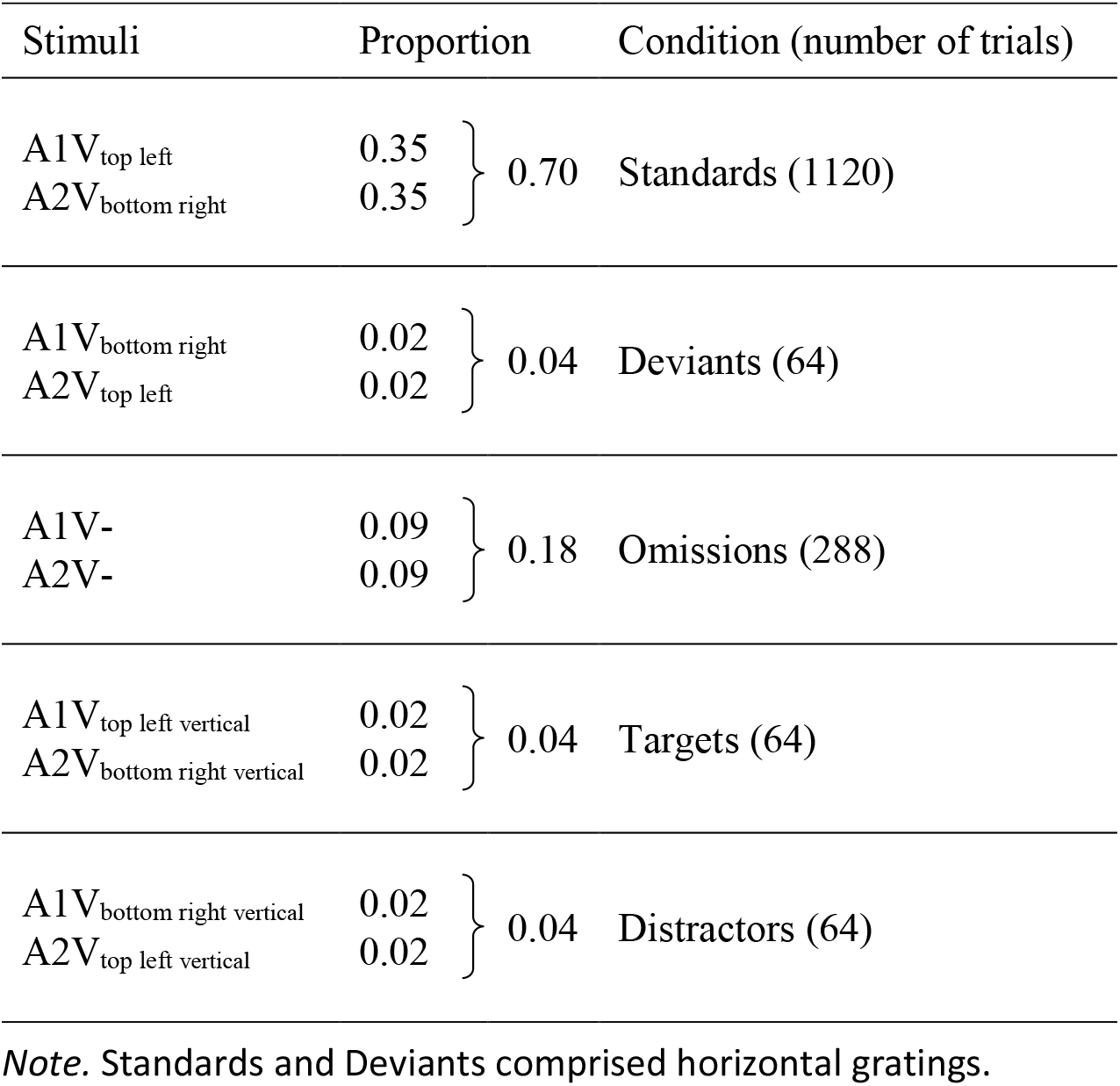
Probabilities for each condition in the crossmodal run.

In order to guarantee crossmodal predictions we made crossmodal combinations (visual location and sound) task relevant. We introduced additional trials (8%) with vertically oriented gratings. Four percent of these trials used the Standard sound-visual location combination (called “Targets”: A1V_top left vertical_, A2V_bottom right vertical_) and the remaining four percent used the Deviant sound-visual location combination (called “Deviants”: A1V_bottom right vertical_, A2V_top left vertical_). Participants had to respond to Target trials only, that is, to respond to trials with vertically oriented gratings if the sound-visual location combination corresponded to the Standard combinations. Participants had to respond within 800 ms after the background switch. Upon a correct response, the fixation cross remained white for another 200 ms; after an incorrect response the fixation cross turned red instead. The next trial started after a random interstimulus interval (ITI), ranging between 1 and 1.4 s (uniform distribution).

#### Unimodal runs

Prior to the crossmodal run, the visual grating stimuli (V_top left_/V_bottom right_) and the sounds (A1/A2) were presented in two separate unimodal baseline runs. The trial structure of the unimodal runs (Figure 1C and 1D) were the same as for crossmodal runs (Figure 1B), except that either only the auditory or only the grating stimuli were presented.

In the visual run, trials started with the presentation of the fixation cross on a black background for 750 ms. Simultaneously with the background switch one of the two possible grating stimuli (V_top left_/V_bottom right_) was presented for 500 ms. Upon stimulus extinction, the background changed back to black. In twenty out of 220 trials (9.1%), vertically oriented gratings were presented: Participants were asked to respond to vertical grating stimuli, irrespectively of stimulus location. In the remaining trials, horizontally oriented gratings were presented (called “Visual only” in the following). As in crossmodal runs, participants had to respond within 800 ms following the background switch. Feedback was provided as in the crossmodal runs. The trial ended with a random ITI, ranging between 1 and 1.4 s (uniform distribution).

In the auditory run, the trials started with the presentation of the fixation cross on a black background and the onset of one of the two auditory stimuli (A1/A2). After 750 ms the background switched from black to gray. The auditory stimulus terminated after an additional 500 ms and the background changed back to black. In twenty of the 220 trials (9.1%), the auditory stimuli contained a gap; these sounds served as targets and required a response. In the remaining trials, the auditory stimuli were presented without the gap (called “Auditory only” in the following). Participants had to respond within 800 ms following the onset of the gap. Feedback was provided as in the crossmodal run. After a random ITI, ranging between 1 and 1.4 s (uniform distribution), the next trial started. Auditory only trials were physically identical to Omissions (A1V-/A2V-) (see Figure 1B and 1D).

#### Procedure

Participants were comfortably seated in a dimly lit room at a 60 cm distance from the screen. The experimental session always began with the unimodal baseline runs. The order of unimodal runs was counterbalanced across participants. Each unimodal run comprised 220 trials. Trials were randomly presented in two blocks of 110 trials each.

After the completion of both unimodal runs, the crossmodal run followed with 16 blocks each consisting of 100 trials. Crossmodal trials were randomized in sets of four subsequent blocks guaranteeing that relative trial probabilities were equally distributed across the whole crossmodal run. Before the crossmodal run started, participants were informed about the likelihood of the frequent combinations of tone and grating location.

#### EEG recording and Preprocessing

The electroencephalogram (EEG) was recorded from 74 Ag/AgCl electrodes positioned according to the 10-10 system (Acharya, Hani, Cheek, Thirumala, & Tsuchida, 2016) and mounted in an elastic cap (EASYCAP GmbH, Herrsching, Germany), with location AFz serving as ground and the left earlobe as reference. The EEG signal was recorded at a sampling rate of 1000 Hz with a BrainAmp DC amplifier (Brain Products GmbH, Gilching, Germany); with a hardware bandpass filter with a passband of 0.0167 to 250 Hz. The electrode impedances were kept below 15kΩ.

Offline EEG data were preprocessed using custom scripts and the EEGLAB toolbox version 14.1.1b (Delorme & Makeig, 2004) for MATLAB version R2015a (MathWorks Inc., Natick, MA, USA). First, we epoched the recordings from -1.75 to 1 s centered at the onset of the background switch and baselined each trial by subtracting the average activity of the -1.75 to 0 ms baseline epoch from each post switch time point. Subsequently, data were low-pass filtered with a finite impulse response filter and an upper cutoff frequency of 110 Hz (6dB cut off at 123.75 Hz, 27.5 Hz transition bandwidth) using the pop_eegfiltnew function in EEGLAB. Electrodes with artifacts (e.g., muscle activity, movement of the electrode, electrode saturation) in more than 15% of the trials were removed (0.18 channels per subject, range: 0 to 2 channels, SD = 0.5) and later substituted by spherical linear interpolation of the three closed neighboring channels.

Next, to remove typical biological (blink, eye movement, muscle, heart) and other (line noise) artifacts we used the Independent Component Analysis (ICA, as implemented by EEGLAB *runica* function) (Lee, Girolami, & Sejnowski, 1999). Components representing artifacts were identified by employing the ICLabel classifier (Pion-Tonachini, Kreutz-Delgado, & Makeig, 2019). This classifier calculates for each Independent Component the probability that it captures brain activity or rather activity related to artifacts. A component was considered as representing an artifact if the probability exceeded 0.8 for one of the artifact categories muscle, eye, heart and line noise. Additionally, based on the scalp topography and power spectrum of eye-movement related Independent Components (Plöchl, Ossandón, & König, 2012) we subsequently added or adapted the labels for components that were not classified as eye components (additional 0.43 components per subject, range: 0 to 3, SD = 0.8). All independent components identified as artifacts were removed (on average 13.55 components per subject, range: 4 to 33, SD = 6.60).

To ensure that participants had perceived the critical time point of visual stimulation, we removed trials in which the participants had blinked during or near the onset of the background switch. This was achieved by removing trials in which the activity of the independent component related to blinks exceeded a threshold of +/-25 standard deviations in a time window of +/-150 ms around the onset of the background switch.

Next, the EEG data were average-referenced and the onset of the event-mark for the background switch was corrected (see Stimuli and Apparatus). Subsequently, we interpolated the previously rejected electrodes. Thereafter we applied a low-pass filter with an upper cut-off at 40 Hz (6 dB cut off at 59.64 Hz, 39.29 Hz transition bandwidth). Epochs were baseline corrected to the average activity between -100 and 0 ms. This baseline was used for statistical analyses. Finally, trials that still contained values exceeding +/-100 µV were removed. Only Standards, Deviants and Omissions of the crossmodal run (see Table 1) and non-target trials of the two unimodal runs (Auditory only and Visual only) were analyzed. Thus, only trials without manual responses and thus motor-related activity were considered. In total, 94.7% of trials of the Auditory only condition, (range = 57% to 100%, *SD* = 8.51), 95.2% of trials of the Visual Only condition (range = 55% to 100%, *SD* = 8.13), 95.1% of trials of the Standard condition (range = 67.14 to 99.82%, *SD* = 8.13), 95.6% of trials in the Deviant condition (range = 68.75 to 100%, *SD* = 6.92) and 94.7% of trials in the Omission condition (range = 67.36 to 100%, *SD* = 6.77) remained for the statistical analysis.

#### Behavioral analysis

The hit rate for targets was defined separately for the crossmodal, visual and auditory only runs as the percentage of correct responses to targets divided by the absolute number of target trials. Correspondingly, the false alarm rate was calculated by dividing the number of responses in non-target trials by the total number of non-target trials (absolute number of trials (Standards, Deviants, Omissions, and Deviants) minus number of target trials).

#### ERP Analysis

Event-related potentials (ERP) were analyzed using the Fieldtrip toolbox (Oostenveld, Fries, Maris, & Schoffelen, 2011) and customized MATLAB scripts.

#### C1 effect

To investigate neural activity at the first cortical stages of visual processing we calculated the C1 effect by subtracting the ERPs elicited by V_bottom right_ visual stimulation from the ERPs elicited by V_top left_ visual stimulation, separately for Standards and Deviants. Moreover we calculated the analogous difference potential for Omissions (crossmodal run) and Auditory only trials (unimodal run), that is we subtracted the ERPs to sounds associated with the V_bottom right_ from the ERPs to tones associated with the V_top left_.

The C1 effect was parameterized by calculating the average voltage of the 50-100 ms post background switch epoch across 20 posterior electrodes (CPz/1/3/5, Pz/1/3/5/7/9, POz/3/7/9, Oz/1/9, Iz, TP7/9, see Figure 2B). The time epoch and the posterior electrode selection were adapted from (Sourav et al., 2018) (see also: (Di Russo et al., 2001). We analyzed the C1 at left hemisphere electrodes that is, at electrodes ipsilateral to the upper and contralateral to the bottom visual stimulation. This decision was based on the previous observations that the C1 wave is largest for upper visual field stimulation at ipsilateral posterior electrodes and larger for lower visual field stimulation at contralateral electrodes (Baumgartner, Graulty, Hillyard, & Pitts, 2018; Di Russo et al., 2001; Kelly, Gomez-Ramirez, & Foxe, 2008). Consequently, the largest C1 effect (ERP difference between upper and lower visual field stimulation) for the two stimulus locations employed in the present experiment is expected for left parieto-occipital recordings. The existence of a C1 effect to Standards, Deviants and Omissions was evaluated with one-tailed t-tests against zero after applying a Bonferroni Correction for multiple testing. To compare the size of the C1 effect across conditions, a one-way repeated measures ANOVA with the repeated measures factor condition (four levels: Standards, Deviants, Omissions and Auditory only) was run. A significant main effect was followed up by post-hoc two-tailed t-tests with Bonferroni correction to compare the C1 effect between conditions.

**Figure 2.**
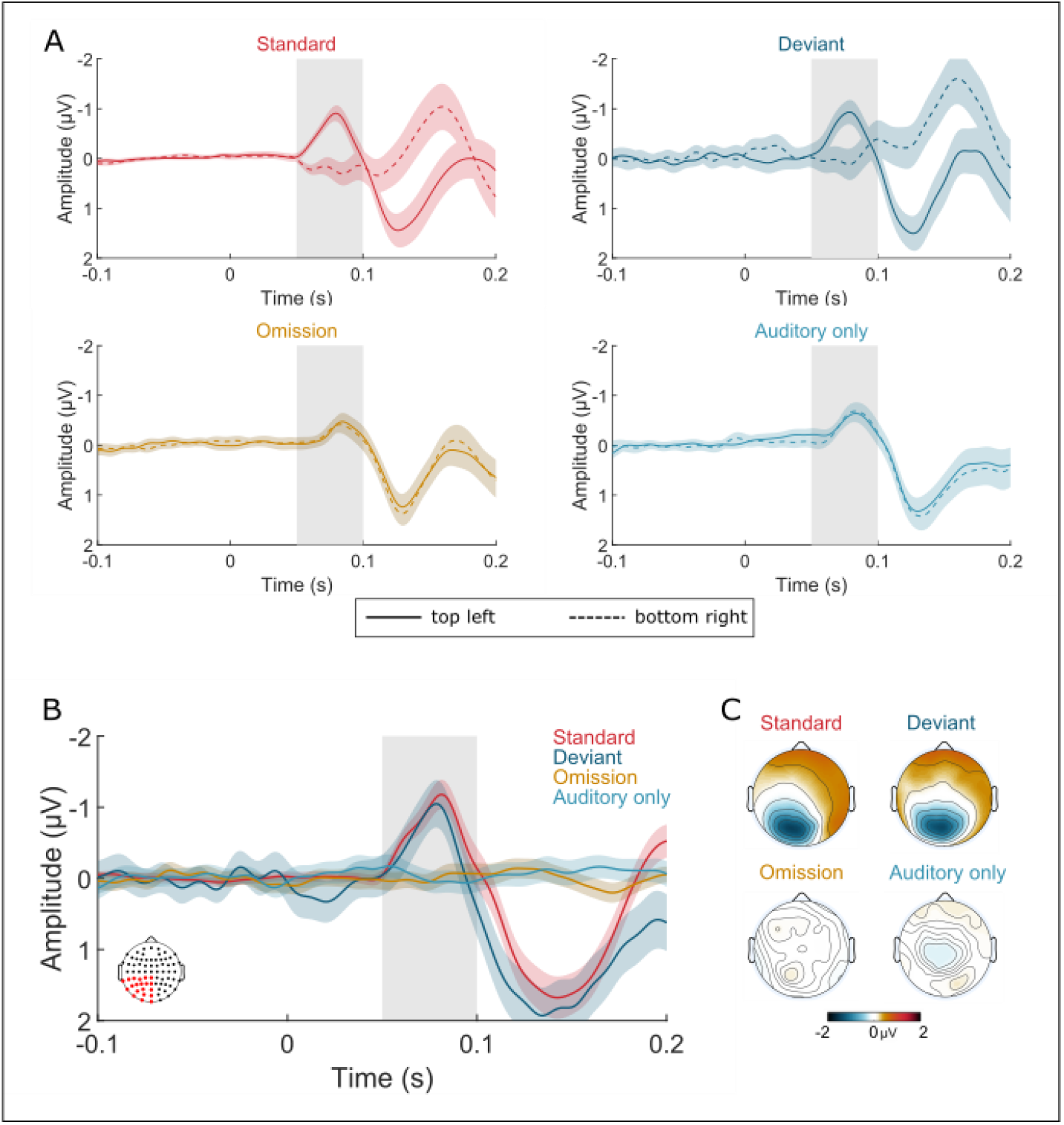
Grand average C1 effects. (A) ERPs elicited by top left (solid line) and bottom right (dashed line) grating stimuli by Standards (left upper panel), Deviants (right upper panel), Omissions (left lower panel) and Auditory only trials (right lower panel). The C1 latency (50-100 ms) is shaded in gray. Error bands represent the standard error of the mean. Time zero is the onset of the background switch. All ERPs are averages across 20 posterior electrodes depicted in the insert in (B). (B) Grand averages of the C1 effect for all four conditions (ERP(V_top left_) minus ERP (V_bottom right_)). (C) ERP topographies displaying the C1 effect separately for all four conditions.

#### Cluster-based permutation analysis

Separate cluster-based permutation tests (Maris & Oostenveld, 2007) as implemented in the Fieldtrip toolbox were employed to evaluate Mismatch and Omission effects following the C1. (See below). All cluster-based permutation tests were run over the 500 ms post background switch time epoch including all scalp electrodes. We defined positive and negative clusters according to the polarity of the difference waves. ERPs to the omission of expected visual stimuli were compared for V_top left_ and V_bottom right_ trials. Any significant ERP difference would indicate a spatially specific omission response. To confirm that the Omission effect was exclusively driven by visual expectations, the same difference was defined for the Auditory only trials and both signals were subsequently compared with an additional cluster based permutation test.

ERPs to Standards vs. Deviants were separately compared for V_top left_ and V _bottom right_ trials. The resulting Mismatch effect indicates a stimulus driven violation of the crossmodal expectation and the difference of both conditions indicates any spatially specific aspect of this the mismatch response. Comparison allowed us to examine electrophysiological activity that was temporally and spatially specific with respect to the original visual prediction.

#### Data and Materials availability

The code for the statistical analyses, figures, and the anonymized, pre-processed data are available at the Research Data Repository of the University of Hamburg (DOI: 10.25592/uhhfdm.11175). Original EEG datasets are available upon request from the corresponding author.

## Results

### Behavior

In both unimodal runs, participants had to respond to rare targets (auditory stimuli with a gap and visual stimuli comprising vertical gratings). In the crossmodal run, target trials were defined as stimuli with vertical gratings combined with the sound of corresponding Standard trials. A high hit rate (auditory only: *M* = 98.38%, *SD* = 3.28; visual only: *M* = 96.88%, *SD* = 3.87; crossmodal: *M* = 84.34%, *SD* = 12.11) and a low false alarm rate (auditory only: *M* = 1.20%, *SD* = 4.60; visual only: *M* = 0.30%, *SD* = 0.48; crossmodal: *M* = 1.00%, SD = 0.98) in each run indicated that participants were engaged and able to perform the tasks successfully.

### C1 effects

To investigate whether visual predictions modulate neural activity at the first stages of visual processing we tested whether a significant C1 effect was obtained for Standards, Deviants, Omissions and Auditory only trials. A significant C1 effect was found for Standards (*M* = -0.67 µV, *SD* = 0.75; *t*_(39)_ = -5.64, *p* < .001) and for Deviants (*M* = -0.47 µV, *SD* = 1.07; *t*_(39)_ = -2.80, *p* = .004; Figure 2A) but neither for Omissions (*M* = - 0.01 µV, *SD* = 0.40; *t*_(39)_ = -0.01, *p* = .46) nor for Auditory only trials (*M* = -0.02 µV, *SD* = 0.46; *t*_(39)_ = 0.05, *p* = .52; Figure 2A). A repeated measures ANOVA with the within-subjects factor condition (Standards, Deviants, Omissions, Auditory only) was significant (*F*_(3, 36)_ = 8.53, *p* < .001; Figure 2B and 2C). The C1 effects for Standards and Deviants was larger compared to both Omissions (*t*_(39)_ = -5.64, *p* < .001; *t*_(39)_ = - 2.82, *p* = .007) and Auditory only trials (*t*_(39)_ = -4.54, *p* < .001; *t*_(39)_ = -2.66, *p* = .011). By contrast, the C1 effect was indistinguishable for on the one hand Omissions and Auditory only trials (*t*_(39)_ = 0.13, *p* = .90) and on the other hand Standards and Deviants (*t*_(39)_ = -1.53, *p* = .13).

### Omission effect

In order to investigate the time course and possible spatial specificity of visual omission response, we compared the ERPs to Omissions to V_top left_ vs. Omissions to V_bottom right_. A significant spatially specific Omission effect started around 230 ms with a negative polarity over the left and a positive polarity over the right hemisphere (positive cluster: from 235 to 500 ms, *p* < .001; negative cluster: from 245 to 500 ms, *p* = .002; Figure 3). In contrast, the corresponding comparison of ERPs to Auditory only trials did not reveal any significant effect (*p* = .15 for the first positive, and *p* = .16 for the first negative cluster). Hence, the Omission effect was significantly larger for the crossmodal than for the Auditory only run (positive cluster: from 223 to 500 ms, *p* = .01; negative cluster: from 238 to 500 ms, *p* = .01; Figure S1).

**Figure 3.**
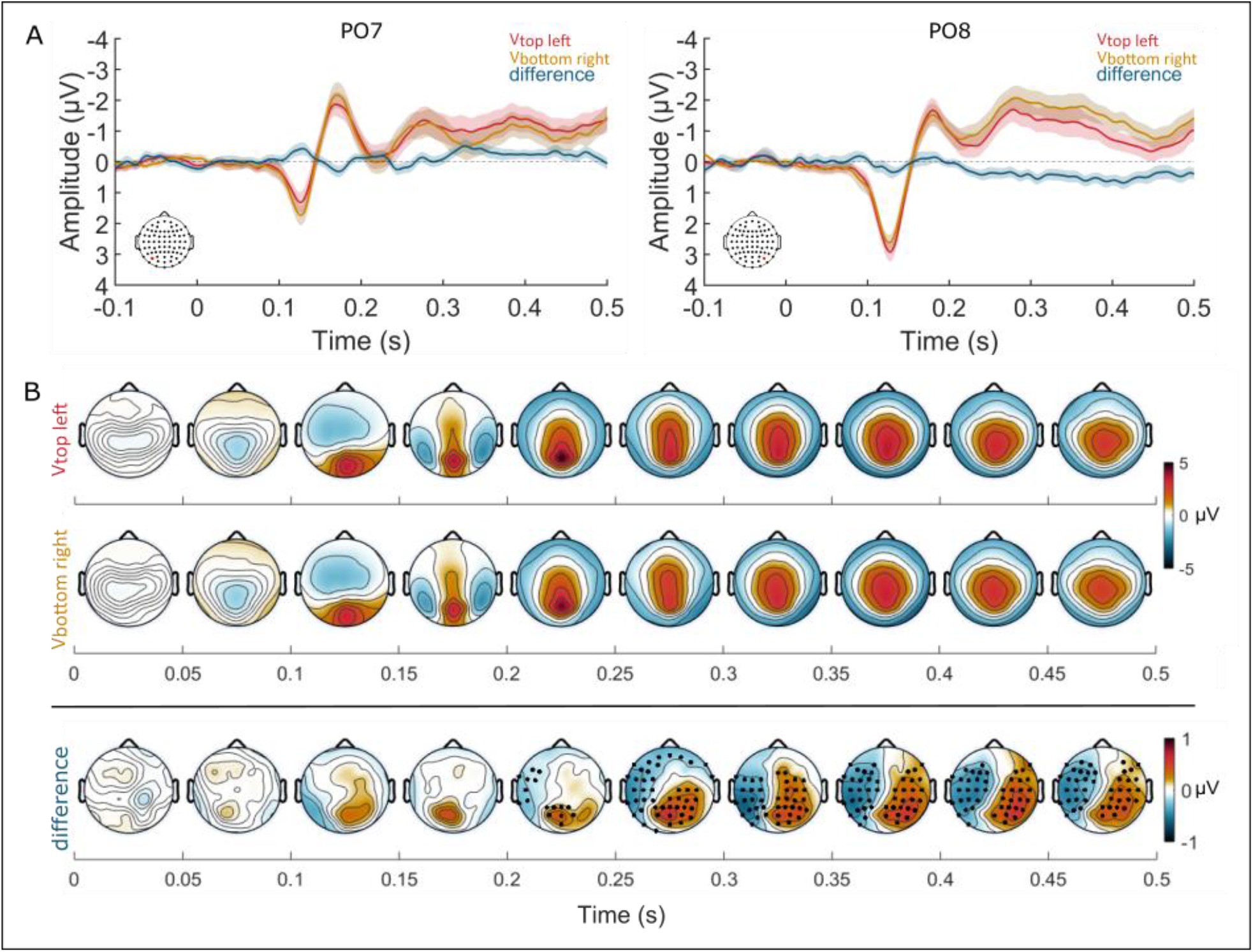
Grand averages of the ERPs to Omissions V_top left_ and V_bottom right_ trials. (A) Omission ERPs to V_top left_ (red) and V_bottom right_ (yellow) locations and the difference ERP (ERP(Omission V_top left_) minus ERP(Omission V_bottom right_)), blue) at electrodes PO7 (left) and PO8 (right) (see red marked electrodes in the schematic drawing of the electrode montage). Error bands represent the standard error of the mean. Time zero is the moment of the background switch. (B) ERP topographies for Omissions to V_top left_ (first row) and V_bottom right_ (second row) location as well as the difference ERP of both conditions (ERP(V_top left_) minus ERP(V_bottom right_); last row) for the post background switch time epoch 0 to 500 ms in 50 ms steps. Stars in the ERP topographies in the last row indicate electrodes that survived tests for multiple comparisons (p < .05).

### Mismatch effects

To investigate Mismatch effects we compared ERPs to Deviants and Standards separately for V_top left_ and V_bottom right_ trials. A significant difference wave was found for the V_top left_ condition over the fronto-central and posterior scalp starting at about 140 ms after the onset of the background switch (positive cluster: from 142 to 500 ms, *p* < .001; first negative cluster: from 176 to 379 ms, *p* < .001; second negative cluster: from 385 to 500 ms, *p* = .005; Figure S2). Corresponding results were obtained for the V_bottom right_ condition (positive cluster: from 142 to 454 ms, *p* < .001; first negative cluster: from 143 to 351 ms, *p* = .002; second negative cluster: from 339 to 500 ms, *p* = .018; Figure S3). Visual stimuli at unexpected location elicited larger ERPs than visual stimuli at the expected location. To investigate the spatially specific aspects of the Mismatch effects we compared the difference ERPs to V_top left_ and V_bottom right_. A significant spatially specific Mismatch response started after around 170 ms after stimulus onset (positive cluster: from 177 to 384 ms, *p* = .006, negative cluster: from 178 to 286 ms, *p* = .021) with a positive polarity over the left hemisphere and a negative polarity over the right hemisphere (Figure 4).

**Figure 4.**
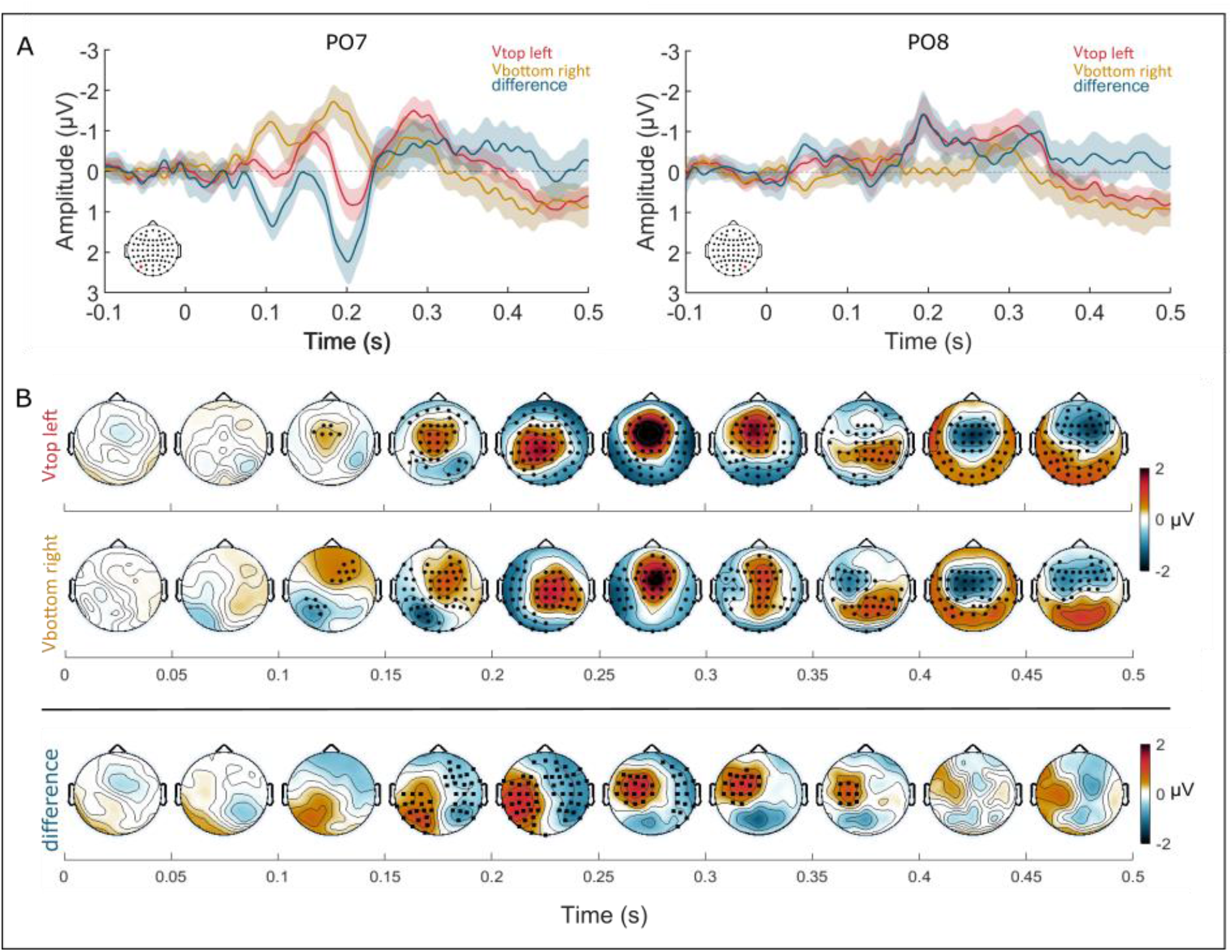
Grand averages of the ERPs for the Mismatch effect V_top left_ and V_bottom right_. (A) ERPs to Mismatch effect V_top left_ (red) and V_bottom right_ (yellow) and the difference ERP (blue) derived by subtracting ERPs to V_top left_ Mismatch from ERPs to V_top left_ Mismatch at electrodes PO7 (left) and PO8 (right) (see red marked electrodes in the schematic drawing of the electrode montage). Error bands represent the standard error of the mean. Time zero is the moment of the background switch. (B) ERP topographies display the ERPs to V_top left_ Mismatch (first row) and V_bottom right_ Mismatch (second row) and the difference of ERPs to V_top left_ Mismatch minus ERPs to V_bottom right_ Mismatch (last row) for the post background switch time epoch 0 to 500 ms in 50 ms steps. Stars in the ERP topographies indicate electrodes that survived tests for multiple comparisons (p < .05).

## Discussion

Previous studies have demonstrated stimulus specific neural activity to omissions of expected visual stimuli in early visual cortex. Yet the precise timing of early visual cortex activity induced by crossmodal predictions remained elusive. Here we implemented a novel crossmodal omission paradigm that allowed testing whether crossmodally visual predictions cause activity in early visual cortex that mimics visual stimulus driven activity.

Two different auditory stimuli were associated with visual grating stimuli at two distinct locations (the top left quadrant and the bottom right quadrant). These two grating stimuli were expected to elicit an event-related potential (ERP) of different polarity between 50 and 100 ms, known as the C1 effect, which is considered to indicate early, likely primary visual cortex activity (Di Russo et al., 2001). If crossmodal visual predictions mimic the processing of real visual stimuli, we would expect to find a C1 effect in trials in which the sound predicted the visual stimulus but the latter was omitted. However, we did not observe a C1 effect to omissions. The first spatially specific omission responses emerged not before 230 ms. Additionally we observed a spatially specific mismatch effect to Deviants starting at about 170 ms.

Functional magnetic resonance imaging (fMRI) studies have convincingly demonstrated that expected but omitted visual stimuli can cause stimulus specific activity in early visual cortex (Kok et al., 2014; Muckli et al., 2015). Moreover, previous experiments employing crossmodal priming have reported a modulation of visual ERPs by preceding sounds (Störmer, McDonald, & Hillyard, 2009). However, the precise timing of visual spatial prediction effects has been unknown. One reason was that fMRI measures do not have the temporal resolution to distinguish activity within the first 500 ms. Another reason was that the employed ERP methods did not use the stimulation protocols necessary to dissociate striate from extrastriate processing or stimulus related processing (e.g. of the auditory cues) from effects of visual predictions. Neural activity to highly expected but omitted visual stimuli are considered to reflect the “pure” visual prediction effect (Egner, Monti, & Summerfield, 2010). The majority of electrophysiological studies in humans have investigated omission responses in the auditory domain (Walsh, McGovern, Clark, & O’Connell, 2020). For example, (Bendixen et al., 2009) reported a high similarity of early (<100 ms) ERPs to omissions and to veridical stimuli. However, (Bendixen et al., 2009) experiment and other previous studies (SanMiguel, Saupe, & Schröger, 2013; Todorovic et al., 2011; Todorovic & Lange, 2012; Wacongne et al., 2011)

The present study implemented a stimulation protocol that unambiguously allowed us to assess the first neural response in visual cortex by investigating the existence of a C1 effect to real expected (and unexpected) visual stimuli and to omissions. While a reliable C1 effect was obtained for real visual stimuli (Standards and Deviants), Omissions did not elicit a C1 effect. Often it was argued that omission responses might not elicit short living early sensory ERPs as the C1 due to the temporal uncertainty associated with omissions. In the present study, we excluded this alternative account by specially marking the time point at which visual stimulation was to be expected: A background switch occurred simultaneously with the presentation of the grating stimulus. Thus, we think that we provided strong evidence in humans for the assumption that the neural mechanisms of early visual cortex activity for visual predictions are distinct from those of processing veridical stimuli (see (Klink, Dagnino, Gariel-Mathis, & Roelfsema, 2017) for non-human primate data; (Keller & Mrsic-Flogel, 2018) for a mechanistic discussion). However, a lateralized and thus spatially specific omission response emerged later, that is at about 230 ms after the expected onset of the visual stimulus. First, this latency of an omission response in the present study concurs with previously observed onsets of omission responses (Ford, Roth, & Kopell, 1976; Klinke, Fruhstorfer, & Finkenzeller, 1968; Simson, Vaughan Jr, & Walter, 1976). Since in contrast to most of the previous studies we had employed lateralized visual stimuli we were able to test whether visual predictions are spatial specific that is, lateralized. In fact, they were. Here it has to be noticed that the ERPs in Omission trials comprised earlier deflections too. However, these were likely elicited by the background switch. Our design did not allow dissociating background switch related neural responses from spatially unspecific omission responses. The lateralized omission response observed in the present study supports the idea that omission responses preserve feature-specific information about the expected stimulus (Demarchi, Sanchez, & Weisz, 2019) and thus collaborates previous fMRI studies (Kok et al., 2014; Muckli et al., 2015). The new findings add that feature-specific response are restricted to late (>200 ms) time epochs and might be associated with a (negative) error signal in visual cortex. Thus, we concluded that visual prediction effects in visual cortex are stimulus specific, but the timing of crossmodally induced top-down activation does not mirror the timing of visual bottom-up activation and thus the neural mechanisms of bottom-up and top-down driven neural activity in early visual cortex are distinct.

A recent ultra-high field fMRI study (Aitken et al., 2020) aimed at precisely characterizing omission responses to expected grating stimuli of different orientations. The authors found a feature-specific omission response in the deep layers 5 and 6 of V1. They discuss their results as evidence for an isolated top-down generated signal analogous to what had been found during mental imagery (Bergmann, Morgan, & Muckli, 2019) and visual working memory tasks (van Kerkoerle, Self, & Roelfsema, 2017). The present results add to the layer specific brain imaging results the precise timing of the stimulus specific omission response: The late onset clearly dissociates top-down activity from stimulus-driven activity in V1 as suggested based on non-human animal work (Keller & Mrsic-Flogel, 2018; Klink et al., 2017).

If visual predictions elicit lateralized neural responses not before 230 ms in the absence of sensory-driven activity, the question arises of whether top-down effects might modulate stimulus-driven activity at an early point in time. (Klink et al., 2017) demonstrated that feedback activity is not capable of generating similar excitatory activity in V1 as feedforward activity from the thalamus. Moreover (Kok, Bains, van Mourik, Norris, & Lange, 2016) reported that while top-down activity predominantly modulated deeper layers, bottom-up activity elicited activity in all cortical layers. Here we investigate effects of crossmodal visual predictions on the processing of visual stimuli by comparing ERPs to stimuli that mismatched vs. matched the crossmodal visual prediction. First at all, although Deviants elicited a C1 effect this effect was not modulated by the visual prediction. In fact, this finding is consistent with a large body of literature failing to find evidence for a modulation of the first cortical response of visual processing by top-down manipulations (Alilović, Timmermans, Reteig, van Gaal, & Slagter, 2019; Baumgartner et al., 2018; Di Russo et al., 2012; Di Russo, Martínez, & Hillyard, 2003; Roelfsema, Lamme, & Spekreijse, 1998; Supèr, Spekreijse, & Lamme, 2001). For example, (Alilović et al., 2019) simultaneously manipulated visual expectations and spatial attention by presenting a predictive word cue at the beginning of each trial (left/right/neutral) and a color cue indicating the side participants had to attend, respectively. In line with our results, they did not observe a C1 modulation related to prediction or to spatial attention (for the seminal work on this topic see (Martinez et al., 1999). Thus, the current and previous evidences converges to the conclusion that the C1 effect indicating retinotopically organized processing between 50-100 ms is not subject of top-down, including crossmodal control. To investigate when in the visual processing pathway visual predictions modulate neural activity evoked by visual stimuli we looked at ERPs following the C1 time range. ERPs to Deviants compared to Standards started to differ after around 140 ms post stimulus presentation, that is, in the time epoch of the N1 of the visual ERPs. The Mismatch effect started to differ for the two visual locations with a latency of about 170 ms. We observed a more negative going potential (mismatch effect) over the contralateral hemisphere. The here observed mismatch effect is reminiscent of a visual mismatch negativity (vMMN) (Sulykos & Czigler, 2011). The vMMN response has been interpreted as reflecting an error signal elicited by visual stimuli that violated the visual expectation (for a review see: (Stefanics, Kremláček, & Czigler, 2014). This error signal was localized to extrastriate visual cortex by (Kimura, Ohira, & Schröger, 2010). Thus, we consider that the spatially specific mismatch response of the present study similarly reflects an error signal associated with extrastriate cortex. (Keller & Mrsic-Flogel, 2018) distinguished two types of error signals for which they postulated different neural mechanisms: First, a negative error signal which is supposed to be generated if the stimulation undershoots the expected activation. Second, a positive error signal to a stimulation that overshoots the expected input. In this terminology, omission trials would trigger negative error signals while unexpected stimuli, such as Deviants in the present study, would be assumed to predominantly generate positive error signals. In the framework of (Press, Kok, & Yon, 2020) Deviants were unexpected but strong events (they comprise a grating stimulus), while Omissions may be considered as unexpected but weak events (not containing a task relevant visual event). Only the processing of unexpected strong events is assumed to be upregulated, possibly reflected in the N1 effect. At a later processing stages, (Press et al., 2020) suggest that the result of the comparison of input and output emerges. Thus, we would conclude that positive error signals are generated to behavioral relevant stimuli earlier to trigger quick model updating, while negative error signals might affect particular the later processing stages.

In sum, the present study demonstrates that cross-modally controlled early visual cortex activity does not follow the same time course as bottom-up driven activity, that is, it does not go together with an retinotopically specific early visual cortex activity between 50 and 100 ms. If visual predictions are wrong, they seem to be associated with spatially specific late (negative) error signals (>200 ms) in the absence of the expected visual simulation and a (positive) error signals (>100 ms) if an unexpected visual event occurs. The modulatory effect of crossmodal visual predictions on visual cortex activity thus seems to be quick and precise enough to modulate visual perception including object recognition (Williams, Markov, Tiurina, & Störmer, 2022).

## Acknowledgments

The authors are grateful to Deniz Främke who helped with data collection.

## Reference List

Acharya, J. N., Hani, A. J., Cheek, J., Thirumala, P., & Tsuchida, T. N. (2016). American clinical neurophysiology society guideline 2: Guidelines for standard electrode position nomenclature. The Neurodiagnostic Journal, 56(4), 245–252.

Aitken, F., Menelaou, G., Warrington, O., Koolschijn, R. S., Corbin, N., Callaghan, M. F., & Kok, P. (2020). Prior expectations evoke stimulus-specific activity in the deep layers of the primary visual cortex. PLoS biology, 18(12), e3001023.

Alilović, J., Timmermans, B., Reteig, L. C., van Gaal, S., & Slagter, H. A. (2019). No evidence that predictions and attention modulate the first feedforward sweep of cortical information processing. Cerebral Cortex, 29(5), 2261–2278.

Alink, A., Schwiedrzik, C. M., Kohler, A., Singer, W., & Muckli, L. (2010). Stimulus predictability reduces responses in primary visual cortex. Journal of Neuroscience, 30(8), 2960–2966.

Banzi, M., & Shiloh, M. (2014). Getting started with Arduino: The open source electronics prototyping platform: Maker Media, Inc.

Bastos, A. M., Usrey, W. M., Adams, R. A., Mangun, G. R., Fries, P., & Friston, K. J. (2012). Canonical microcircuits for predictive coding. Neuron, 76(4), 695–711.

Baumgartner, H. M., Graulty, C. J., Hillyard, S. A., & Pitts, M. A. (2018). Does spatial attention modulate the earliest component of the visual evoked potential? Cognitive neuroscience, 9(1-2), 4–19.

Bendixen, A., Schröger, E., & Winkler, I. (2009). I heard that coming: Event-related potential evidence for stimulus-driven prediction in the auditory system. Journal of Neuroscience, 29(26), 8447–8451.

Bergmann, J., Morgan, A. T., & Muckli, L. (2019). Two distinct feedback codes in V1 for ‘real’and ‘imaginary’internal experiences. bioRxiv, 664870.

Brainard, D. H. (1997). The psychophysics toolbox. Spatial vision, 10(4), 433–436.

de Lange, F. P., Heilbron, M., & Kok, P. (2018). How do expectations shape perception? Trends in cognitive sciences, 22(9), 764–779.

de Lange, F. P., Rahnev, D. A., Donner, T. H., & Lau, H. (2013). Prestimulus oscillatory activity over motor cortex reflects perceptual expectations. Journal of Neuroscience, 33(4), 1400–1410.

Delorme, A., & Makeig, S. (2004). EEGLAB: An open source toolbox for analysis of single-trial EEG dynamics including independent component analysis. Journal of neuroscience methods, 134(1), 9–21.

Demarchi, G., Sanchez, G., & Weisz, N. (2019). Automatic and feature-specific prediction-related neural activity in the human auditory system. Nature communications, 10(1), 1–11.

Den Ouden, H. E. M., Friston, K. J., Daw, N. D., McIntosh, A. R., & Stephan, K. E. (2008). A dual role for prediction error in associative learning. Cerebral Cortex, 19(5), 1175–1185.

Di Russo, F., Martínez, A., & Hillyard, S. A. (2003). Source analysis of event-related cortical activity during visuo-spatial attention. Cerebral Cortex, 13(5), 486–499.

Di Russo, F., Martínez, A., Sereno, M. I., Pitzalis, S., & Hillyard, S. A. (2001). Cortical sources of the early components of the visual evoked potential. Human brain mapping, 15(2), 95–111.

Di Russo, F., Stella, A., Spitoni, G., Strappini, F., Sdoia, S., Galati, G., et al. (2012). Spatiotemporal brain mapping of spatial attention effects on pattern-reversal ERPs. Human brain mapping, 33(6), 1334– 1351.

Egner, T., Monti, J. M., & Summerfield, C. (2010). Expectation and surprise determine neural population responses in the ventral visual stream. Journal of Neuroscience, 30(49), 16601–16608.

Feng, W., Störmer, V. S., Martinez, A., McDonald, J. J., & Hillyard, S. A. (2014). Sounds activate visual cortex and improve visual discrimination. Journal of Neuroscience, 34(29), 9817–9824.

Ford, J. M., Roth, W. T., & Kopell, B. S. (1976). Attention effects on auditory evoked potentials to infrequent events. Biological Psychology, 4(1), 65–77.

Keller, G. B., & Mrsic-Flogel, T. D. (2018). Predictive processing: A canonical cortical computation. Neuron, 100(2), 424–435.

Kelly, S. P., Gomez-Ramirez, M., & Foxe, J. J. (2008). Spatial attention modulates initial afferent activity in human primary visual cortex. Cerebral Cortex, 18(11), 2629–2636.

Kimura, M., Ohira, H., & Schröger, E. (2010). Localizing sensory and cognitive systems for pre-attentive visual deviance detection: an sLORETA analysis of the data of Kimura et al.(2009). Neuroscience letters, 485(3), 198–203.

Kleiner, M., Brainard, D., & Pelli, D. (2007). What’s new in Psychtoolbox-3? Perception 36 ECVP Abstract Supplement. PLOS ONE.

Klink, P. C., Dagnino, B., Gariel-Mathis, M.-A., & Roelfsema, P. R. (2017). Distinct feedforward and feedback effects of microstimulation in visual cortex reveal neural mechanisms of texture segregation. Neuron, 95(1), 209-220. e3.

Klinke, R., Fruhstorfer, H., & Finkenzeller, P. (1968). Evoked responses as a function of external and stored information. Electroencephalography and Clinical Neurophysiology, 25(2), 119–122.

Kok, P., Bains, L. J., van Mourik, T., Norris, D. G., & Lange, F. P. de (2016). Selective activation of the deep layers of the human primary visual cortex by top-down feedback. Current Biology, 26(3), 371– 376.

Kok, P., Failing, M. F., & Lange, F. P. de (2014). Prior expectations evoke stimulus templates in the primary visual cortex. Journal of Cognitive Neuroscience, 26(7), 1546–1554.

Lee, T.-W., Girolami, M., & Sejnowski, T. J. (1999). Independent component analysis using an extended infomax algorithm for mixed subgaussian and supergaussian sources. Neural computation, 11(2), 417– 441.

Maris, E., & Oostenveld, R. (2007). Nonparametric statistical testing of EEG-and MEG-data. Journal of neuroscience methods, 164(1), 177–190.

Martinez, A., Anllo-Vento, L., Sereno, M. I., Frank, L. R., Buxton, R. B., Dubowitz, D. J., et al. (1999). Involvement of striate and extrastriate visual cortical areas in spatial attention. Nature neuroscience, 2(4), 364–369.

Muckli, L., Martino, F. de, Vizioli, L., Petro, L. S., Smith, F. W., Ugurbil, K., et al. (2015). Contextual feedback to superficial layers of V1. Current Biology, 25(20), 2690–2695.

Oostenveld, R., Fries, P., Maris, E., & Schoffelen, J.-M. (2011). FieldTrip: Open source software for advanced analysis of MEG, EEG, and invasive electrophysiological data. Computational intelligence and neuroscience, 2011.

Pion-Tonachini, L., Kreutz-Delgado, K., & Makeig, S. (2019). ICLabel: An automated electroencephalographic independent component classifier, dataset, and website. NeuroImage, 198, 181–197.

Plöchl, M., Ossandón, J. P., & König, P. (2012). Combining EEG and eye tracking: identification, characterization, and correction of eye movement artifacts in electroencephalographic data. Frontiers in human neuroscience, 6, 278.

Press, C., Kok, P., & Yon, D. (2020). The perceptual prediction paradox. Trends in cognitive sciences, 24(1), 13–24.

Puri, A. M., & Wojciulik, E. (2008). Expectation both helps and hinders object perception. Vision research, 48(4), 589–597.

Puri, A. M., Wojciulik, E., & Ranganath, C. (2009). Category expectation modulates baseline and stimulus-evoked activity in human inferotemporal cortex. Brain research, 1301, 89–99.

Roelfsema, P. R., Lamme, V. af, & Spekreijse, H. (1998). Object-based attention in the primary visual cortex of the macaque monkey. Nature, 395(6700), 376–381.

SanMiguel, I., Saupe, K., & Schröger, E. (2013). I know what is missing here: Electrophysiological prediction error signals elicited by omissions of predicted” what” but not” when”. Frontiers in human neuroscience, 7, 407.

Simson, R., Vaughan Jr, H. G., & Walter, R. (1976). The scalp topography of potentials associated with missing visual or auditory stimuli. Electroencephalography and Clinical Neurophysiology, 40(1), 33– 42.

Sourav, S., Bottari, D., Kekunnaya, R., & Röder, B. (2018). Evidence of a retinotopic organization of early visual cortex but impaired extrastriate processing in sight recovery individuals. Journal of vision, 18(3), 22.

Stefanics, G., Kremláček, J., & Czigler, I. (2014). Visual mismatch negativity: A predictive coding view. Frontiers in human neuroscience, 8, 666.

Störmer, V. S., McDonald, J. J., & Hillyard, S. A. (2009). Cross-modal cueing of attention alters appearance and early cortical processing of visual stimuli. Proceedings of the National Academy of Sciences, 106(52), 22456–22461.

Sulykos, I., & Czigler, I. (2011). One plus one is less than two: Visual features elicit non-additive mismatch-related brain activity. Brain research, 1398, 64–71.

Supèr, H., Spekreijse, H., & Lamme, V. af (2001). Two distinct modes of sensory processing observed in monkey primary visual cortex (V1). Nature neuroscience, 4(3), 304–310.

Todorovic, A., & Lange, F. P. de (2012). Repetition suppression and expectation suppression are dissociable in time in early auditory evoked fields. Journal of Neuroscience, 32(39), 13389–13395.

Todorovic, A., van Ede, F., Maris, E., & Lange, F. P. de (2011). Prior expectation mediates neural adaptation to repeated sounds in the auditory cortex: An MEG study. The Journal of neuroscience : the official journal of the Society for Neuroscience, 31(25), 9118–9123.

van Kerkoerle, T., Self, M. W., & Roelfsema, P. R. (2017). Layer-specificity in the effects of attention and working memory on activity in primary visual cortex. Nature communications, 8(1), 1–14.

Wacongne, C., Labyt, E., van Wassenhove, V., Bekinschtein, T., Naccache, L., & Dehaene, S. (2011). Evidence for a hierarchy of predictions and prediction errors in human cortex. Proceedings of the National Academy of Sciences, 108(51), 20754–20759.

Walsh, K. S., McGovern, D. P., Clark, A., & O’Connell, R. G. (2020). Evaluating the neurophysiological evidence for predictive processing as a model of perception. Annals of the New York Academy of Sciences, 1464(1), 242.

Williams, J. R., Markov, Y. A., Tiurina, N. A., & Störmer, V. S. (2022). What You See Is What You Hear: Sounds Alter the Contents of Visual Perception. Psychological science, 09567976221121348.

