## Supplementary material for "Crossmodal visual predictions elicit spatially specific early visual cortex activity but later than real visual stimuli": Figure S1

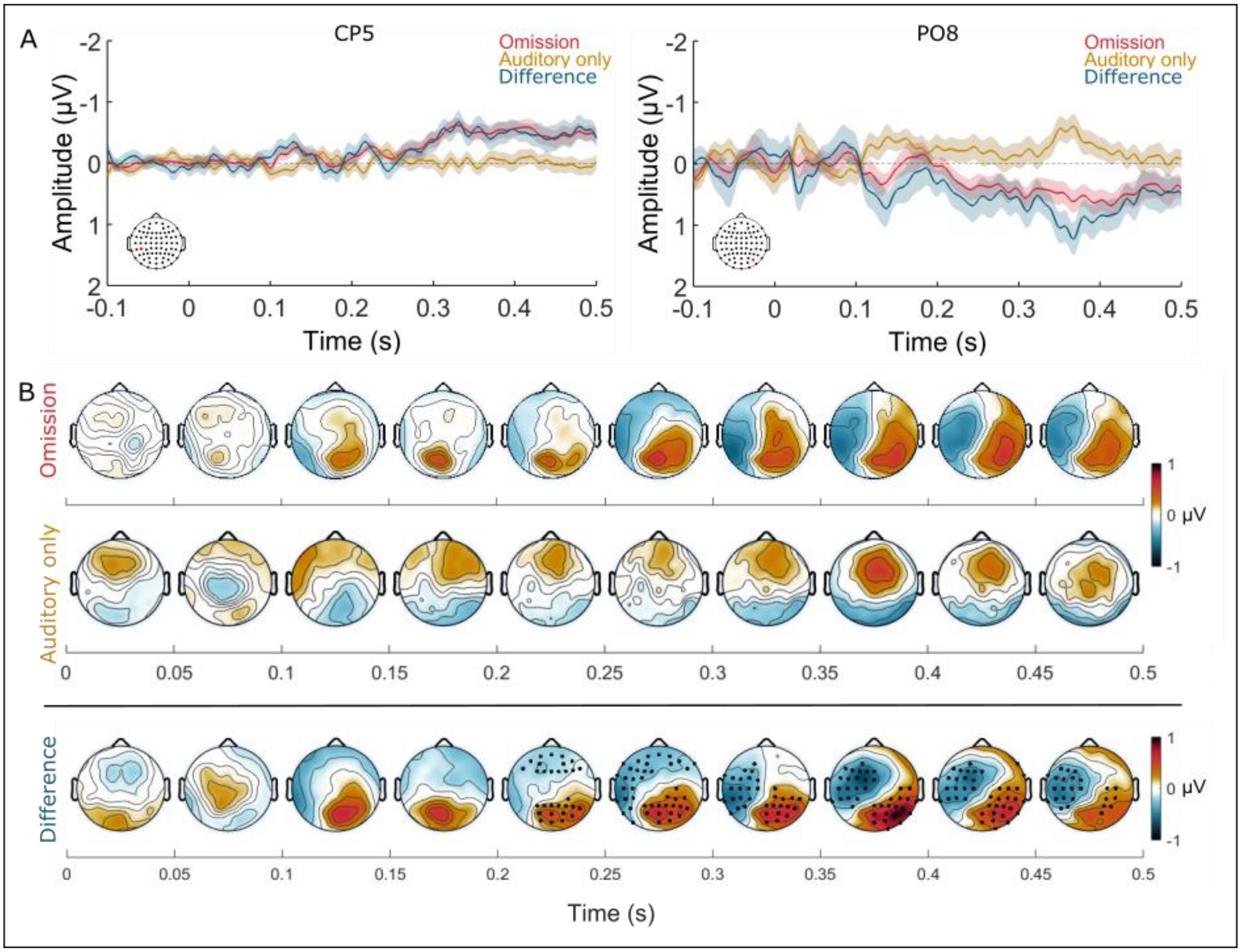


**Figure S1. Grand averages of the ERPs to Omissions and Auditory only.** (A) Difference ERPs to $V_{top\ left}$ Omissions minus $V_{bottom\ right}$ Omissions (red) and difference ERPs to $V_{top\ left}$ Auditory only minus $V_{bottom\ right}$ Auditory only (yellow) and the difference ERP (blue) derived by subtracting the difference ERPs to Auditory only from the difference ERPs for Omissions. ERPs are at electrodes CP5 (left) and PO8 (right) (see red marked electrodes in the schematic drawing of the electrode montage). Error bands represent the standard error of the mean. Time zero is the moment of the background switch. (B) ERP topographies display the difference ERPs for Omissions ($V_{top\ left}$ - $V_{bottom\ right}$; first row) and for Auditory only ($V_{top\ left}$ - $V_{bottom\ right}$; second row) and as well as the difference ERP of both conditions (Omissions($V_{top\ left}$ - $V_{bottom\ right}$) – Auditory only( $V_{top\ left}$ - $V_{bottom\ right}$); last row) for the post background switch time epoch 0 to 500 ms in 50 ms steps. Stars on the ERP topographie in the last row indicate electrodes that survived tests for multiple comparisons ($p < .05$).
