## Supplementary material for "Crossmodal visual predictions elicit spatially specific early visual cortex activity but later than real visual stimuli": Figure S3

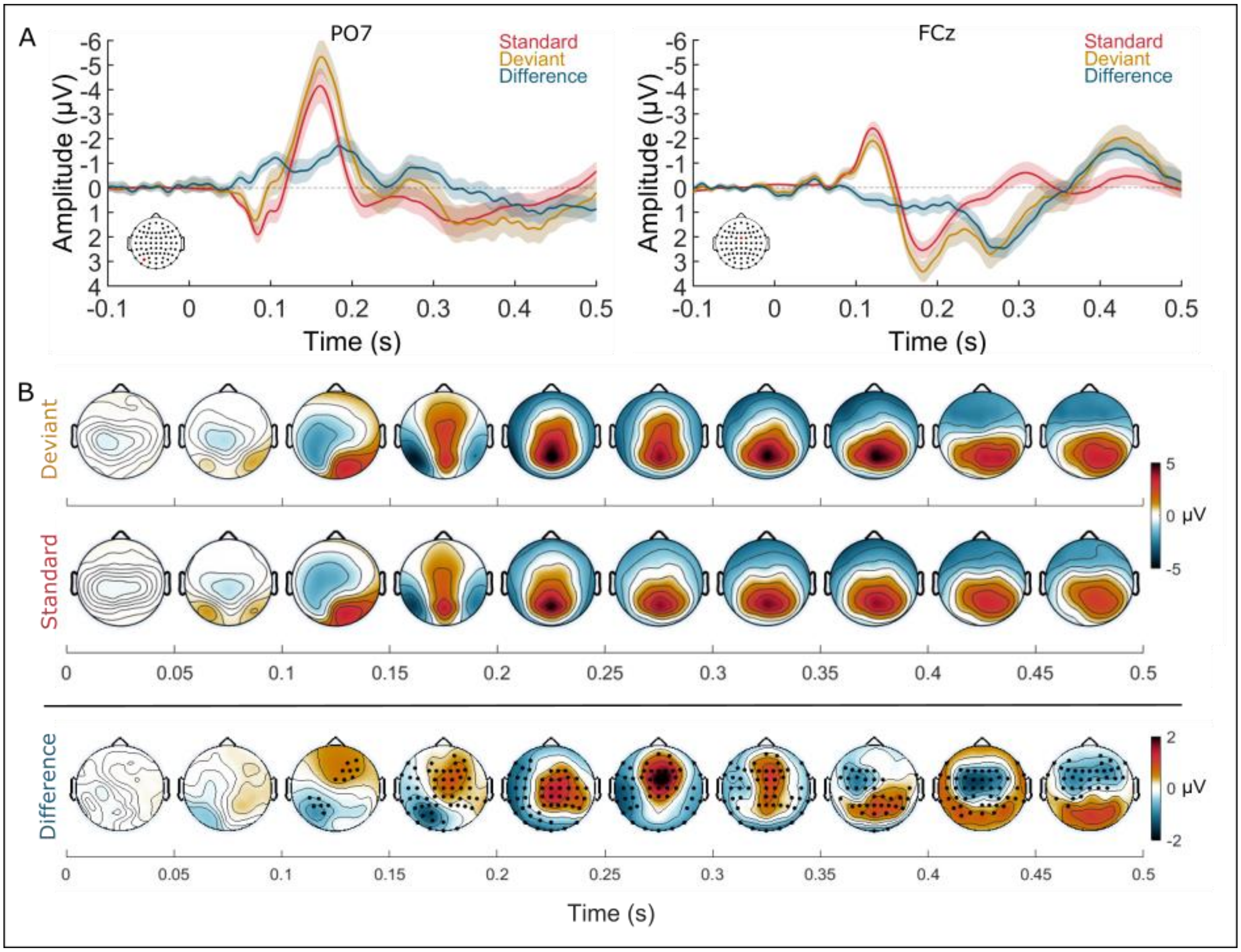


**Figure S3. Grand averages of the ERPs to $V_{bottom\ right}$ Standards and Deviants.** (A) ERPs to $V_{bottom\ right}$ Standards (red) and Deviants (yellow) and the difference ERP (blue) derived by subtracting ERPs to $V_{bottom\ right}$ Deviants from ERPs to $V_{bottom\ right}$ Standards at electrodes PO8 (left) and FCz (right) (see red marked electrodes in the schematic drawing of the electrode montage). Error bands represent the standard error of the mean. Time zero is the moment of the background switch. (B) ERP topographies display the ERPs to $V_{bottom\ right}$ Deviants (first row) and Standards (second row) and the difference of ERPs to $V_{bottom\ right}$ Standards minus ERPs to $V_{bottom\ right}$ Deviants (last row) for the post background switch time epoch 0 to 500 ms in 50 ms steps. Stars in the ERP topographies in the last row indicate electrodes that survived tests for multiple comparisons ($p < .05$).
